# Bad bugs, new drugs: The antimicrobial peptide C14R is active against the ESKAPE pathogens

**DOI:** 10.64898/2025.12.16.694556

**Authors:** Daniel Gruber, Verena Vogel, Jan-Christoph Walter, Sabine Szunerits, Armando Rodríguez, Nico Preising, Ludger Ständker, Carolina Firacative, Barbara Spellerberg, Ann-Kathrin Kissmann, Frank Rosenau

## Abstract

The global rise of antimicrobial resistance among the ESKAPE pathogens represents a major challenge to public health. Here, we report the broad-spectrum antibacterial activity of the synthetic antimicrobial and pore-forming peptide C14R against all six ESKAPE species. Using a radial diffusion assay and resazurin-based viability testing, C14R exhibited potent bactericidal effect with minimum inhibitory concentrations (MICs), defined as the lowest concentration of an antimicrobial agent that completely inhibits visible growth of planktonic microorganisms, ranging from 3.4 μg/mL (*Enterococcus faecium*, vancomycin-resistant) to 45.2 μg/mL (*Klebsiella quasipneumoniae*, ESBL). C14R also inhibited biofilm formation by Gram-positive pathogens, with minimum biofilm inhibitory concentrations (MBICs), referring to the minimal concentration required to prevent the development of biofilms, of 15.0 μg/mL (*Staphylococcus aureus*, MRSA) and 22.0 μg/mL (*E. faecium*, VRE), whereas Gram-negatives biofilms showed higher tolerance. Together, these findings demonstrate that C14R retains high activity against multidrug-resistant ESKAPE strains, highlighting its potential as a lead compound for the development of next-generation antimicrobial drugs to expand the portfolio of available antibiotics and brace health systems against emerging severe infections.

**Author summary:** Antibiotic-resistant infections are a growing threat worldwide. A small group of hospital-associated bacteria is especially problematic because they often evade multiple drugs and cause hard-to-treat infections. In this study, we tested the designed antimicrobial peptide C14R as a novel and effective way to fight these bacteria. Peptides are short protein fragments with the ability to puncture and disrupt microbial membranes. We evaluated C14R against six hospital related priority species (so called ESKAPE pathogens) and measured its ability to stop growth and to limit biofilm formation. C14R killed every species we tested and reduced biofilm of two bacteria. Our findings identify C14R as a promising lead for new treatments, particularly for difficult infections and those involving biofilms.

## Introduction

The Infectious Diseases Society of America (IDSA) reported “Bad Bugs, No Drugs” and its “Call to Action”, which highlighted the threat posed by ESKAPE pathogens (*Enterococcus faecium, Staphylococcus aureus, Klebsiella quasipneumoniae, Acinetobacter baumannii, Pseudomonas aeruginosa, Enterobacter*-species). While their foresight deserves recognition, the persistence of these organisms as a major problem today underscores the urgent need to accelerate discovery and development of new antibacterial drugs (1)(2)(3). Multi-drug-resistant (MDR) bacteria, especially the ESKAPE pathogens are a critical global challenge, causing difficult to treat nosocomial infections (4). They require urgent new solutions for infection control as they often “escape” the effect of commonly used antimicrobial drugs (5)(6). In fact, the World Health Organization (WHO) listed therefore all of the six ESKAPE pathogens in its “2024 Bacterial Priority Pathogen List (WHO BPPL)” underscoring the relevance to find treatment alternatives, whereas rising antimicrobial resistance is not only a problem for third-world-countries but challenges also healthcare in first world countries (7)(8).

The process of “escaping” the effects of the antimicrobial agents relies on multiple resistance mechanisms the microorganisms developed over time. Resistance against vancomycin is frequently observed, as seen in vancomycin-resistant *E. faecalis* (VRE). Vancomycin binds to a precursor involved in peptidoglycan maturation, preventing penicillin-binding proteins (PBPs) from cross-linking lipid II into mature peptidoglycan, thereby weakening the structural integrity of the cell envelope (9)(10)(11). In Methicillin-resistant *S. aureus* (MRSA) the methicillin resistance is mediated by the *mecA* gene, which encodes penicillin-binding protein 2A (PBP2A), characterized by its reduced affinity for β-lactam antibiotics (12). MRSA can compensate the inhibited PBP1 activity by utilizing the “novel” PBP2A, thereby maintaining peptidoglycan cross-linking and enabling growth even in the presence of β-lactam antibiotics such as methicillin (13). *K. pneumoniae* and *Escherichia coli* are frequently found to carry extended-spectrum β-lactamases (ESBLs), making both to clinically relevant strains (14). These enzymes are hydrolyzing the β-lactam ring of a wide range of β-lactam antibiotics including the so-called “third-generation cephalosporins” (15) (16)(17). In the case of *Acinetobacter spp*. MDR is enhanced by horizontal gene transfer of plasmids, transposons and integrons carrying multiple antibiotic genes, as well as by low outer membrane permeability and efflux pumps that reduce drug accumulation (18). *P. aeruginosa* exhibits intrinsic resistance to many antibiotics through low outer membrane porin permeability, active efflux pump systems and chromosomally encoded AmpC β-lactamase, with resistance further enhanced by mutations or acquisition of carbapenemase genes, ultimately leading to MDR strains (2)(19)(20)(21). The antimicrobial peptide C14R is a promising antimicrobial agent, known for its strong, broad-spectrum activity notably against f.e. *P. aeruginosa* by forming pores in their cell membranes (22)(23)(24)(25). In this work, the antimicrobial effect of C14R against different clinically relevant ESKAPE pathogens is evaluated.

## Results

The antimicrobial activity of the synthetic peptide C14R against six clinically relevant ESKAPE pathogens was evaluated using the two-layer radial diffusion assay (Fig 1A+B). Clear inhibition zones were observed for all six strains, indicating that C14R exerts broad-spectrum antibacterial effects. The extent of inhibition varied among species, with *E. faecium* and *P. aeruginosa* showing larger inhibition zones, while *S. aureus* displayed the smallest one.

**Fig 1:**
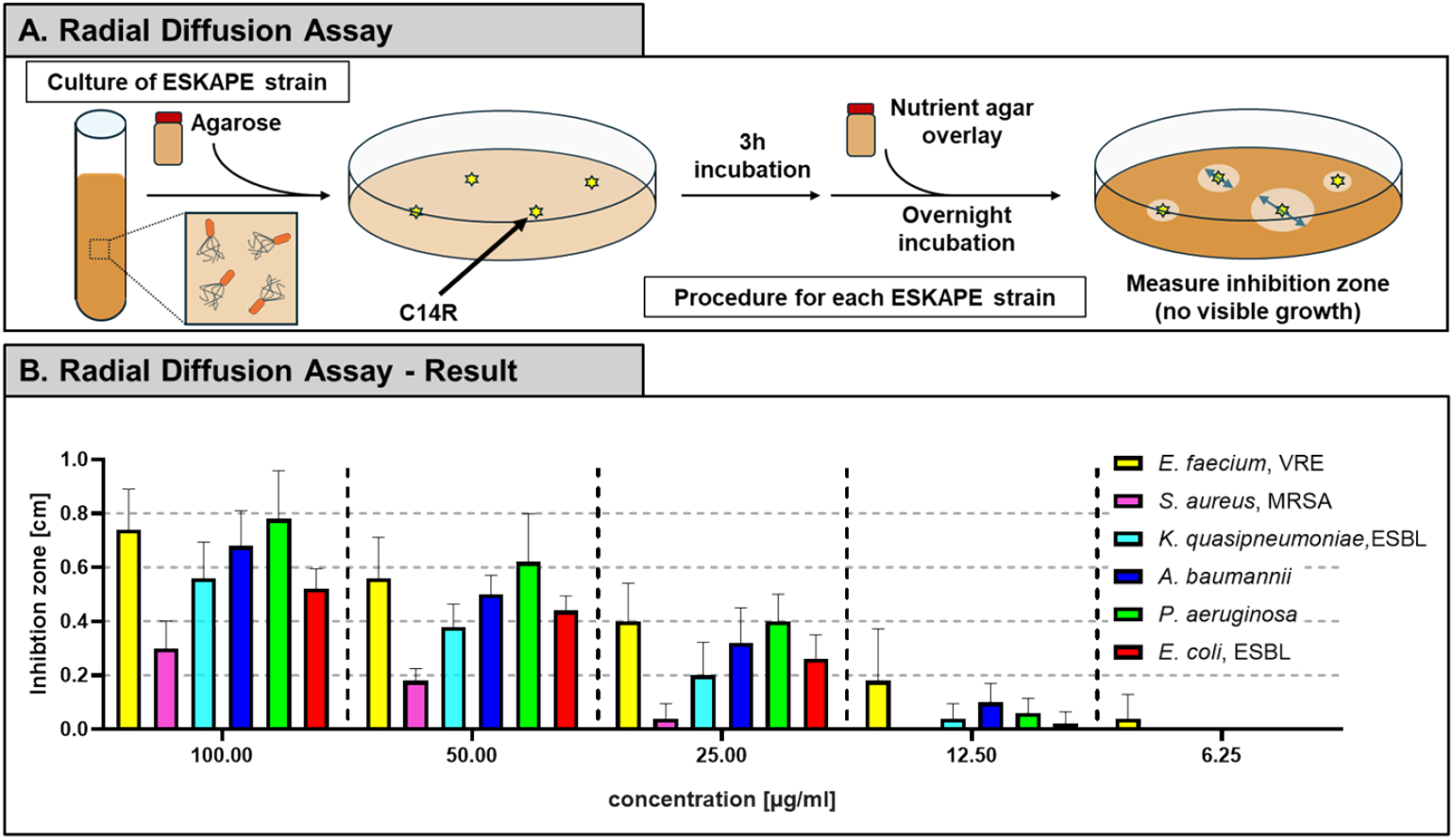
Determination of the antimicrobial activity of C14R using a two-layer radial diffusion assay. (A) Schematic illustration of the two-layer radial diffusion assay. Target bacterial strains were embedded in a first agarose layer and wells were punched into the solidified medium. Wells were filled with serial dilutions of the antimicrobial peptide C14R (100, 50, 25, 12.5 and 6.25μg/mL). After 3 h of pre-incubation at 37°C, a second layer of Trypticase Soy Agar was poured on top. Plates were incubated overnight at 37°C. (B) Measurement of the inhibition zones (cm) for size ESKAPE strains at different C14R concentrations. Data are represented as mean +/- SD, with n = 3 replicates.

To further quantify the antimicrobial potency, resazurin-based viability assays were performed with increasing concentrations of C14R. A clear, concentration-dependent reduction in viability was detected for all pathogens (Fig 2A+2C). The most susceptible strain was *E. faecium* (VRE) with a MIC of 3.44 μg/mL, followed by *S. aureus* (MRSA) with 14.72 μg/mL, *A. baumannii* (23.10 μg/mL) and *E. coli* (ESBL, 24.55 μg/mL). *P. aeruginosa* required slightly higher concentrations for complete inhibition (30.08 μg/mL), while *K. quasipneumoniae* (ESBL), showed the highest tolerance with a MIC of 45.15 μg/mL (Table 1). These data confirm that C14R exhibits broad but differential activity against clinically relevant Gram-positive and Gram-negative pathogens.

**Table 1:**
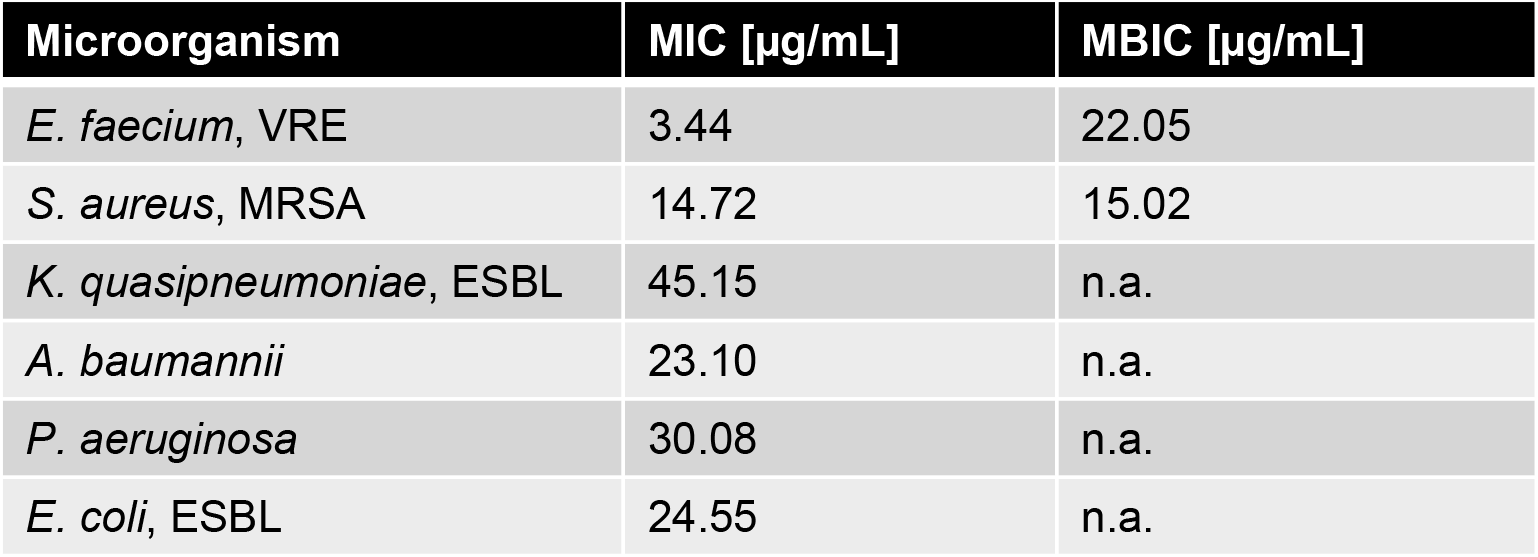
Summary of MIC and MBIC of ESKAPE pathogens (n.a. = not applicable)

| Microorganism | MIC [µg/mL] | MBIC [µg/mL] |
| --- | --- | --- |
| *E. faecium*, VRE | 3.44 | 22.05 |
| *S. aureus*, MRSA | 14.72 | 15.02 |
| *K. quasipneumoniae*, ESBL | 45.15 | n.a. |
| *A. baumannii* | 23.10 | n.a. |
| *P. aeruginosa* | 30.08 | n.a. |
| *E. coli*, ESBL | 24.55 | n.a. |

**Fig 2:**
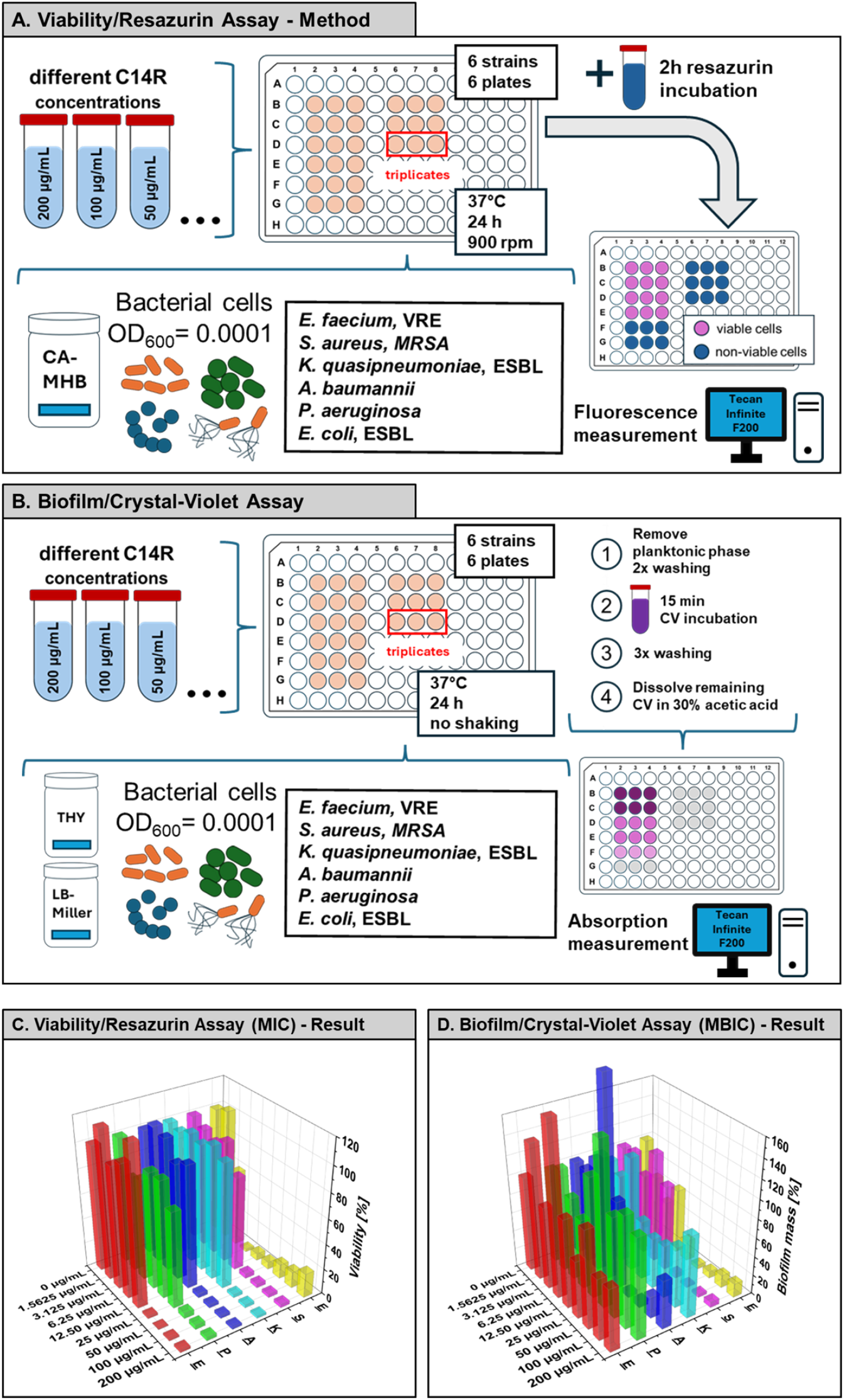
Experimental setup and results of antimicrobial and antibiofilm activity assays for C14R against ESKAPE pathogens. (A) Schematic overview of the resazurin-based viability assay with ESKAPE strains in the presence of C14R (0-200 μg/mL) using cation adjusted Mueller-Hinton Broth (CAMHB) as standard medium for susceptibility testing according to Clinical & Laboratory Standards Institute (CLSI) M100 guidelines (27). (B) Schematic overview of the biofilm/crystal violet assay for measuring the biofilm mass in high nutrient medium, so either Todd-Hewitt (THY) medium or LB-medium supplemented with C14R (0-200 μg/mL). Attached biofilms were stained with crystal violet, washed and solubilized for quantification. (C) Results of the resazurin-based susceptibility assay showing the viability of planktonic ESKAPE cells after 24 h exposure to increasing C14R concentrations (D) Results of the crystal-violet biofilm assay showing relative biofilm formation during a 24 hour exposition to different concentrations of C14R.

Since biofilm formation is a major virulence factor of ESKAPE pathogens (26), the effect of C14R on biofilm development was assessed using the crystal-violet-assay (Fig 2B). In contrast to the MIC assays, a nutrient-rich medium rather than a minimal-medium was used for the MBIC assays to ensure optimal biofilm growth conditions, as biofilm formation in minimal medium was not sufficiently robust. This may account for the higher MBIC compared to the MIC values observed within the same strain. Biofilm biomass decreased in a dose-dependent manner for *E. faecium* and *S. aureus*, with minimum biofilm inhibitory concentration (MBIC) for *E. faecium* of 22.05 μg/mL and *S. aureus* of 15.02 μg/mL (Table 1). For the other ESKAPE pathogens biofilm mass was not or even slightly reduced but not consistently enough therefore no reliable MBIC values could be determined (not determinable, n.d., Table 1). Notably, *S. aureus* biofilms were strongly affected by C14R at intermediate concentrations, whereas *K. quasipneumoniae* was least responsive, reflecting its higher MIC value.

These findings demonstrate that C14R possesses strong antibacterial activity against planktonic and partly against biofilm formation of ESKAPE pathogens, whereas the effect against the planktonic phase was higher against the Gram-positive (*E. faecium, S. aureus*) but also inhibited Gram-negative strains.

In summary, the quantitative analysis of the viability (MIC) and biofilm assays (MBIC) confirmed the activity of C14R against all ESKAPE pathogens, however, with different extends (Table 1). For entries marked as n.a. (not applicable), MBIC values could not be calculated, as MBIC can only be assessed when biofilm formation is reduced to zero, which was not achieved in these cases.

Membrane permeabilization assay first demonstrated that treatment of all ESKAPE pathogens with their strain-specific MICs for C14R induced pore formation, reflected by increased SYTOX Green fluorescence signals across all species (Fig. 3A+C). Correspondingly, colony forming units (CFU) enumeration confirmed a marked loss of viability, with strongly reduced or mostly no colony counts compared to untreated controls (Fig. 3B+C), thereby validating that the determined MIC is biologically active.

**Fig 3:**
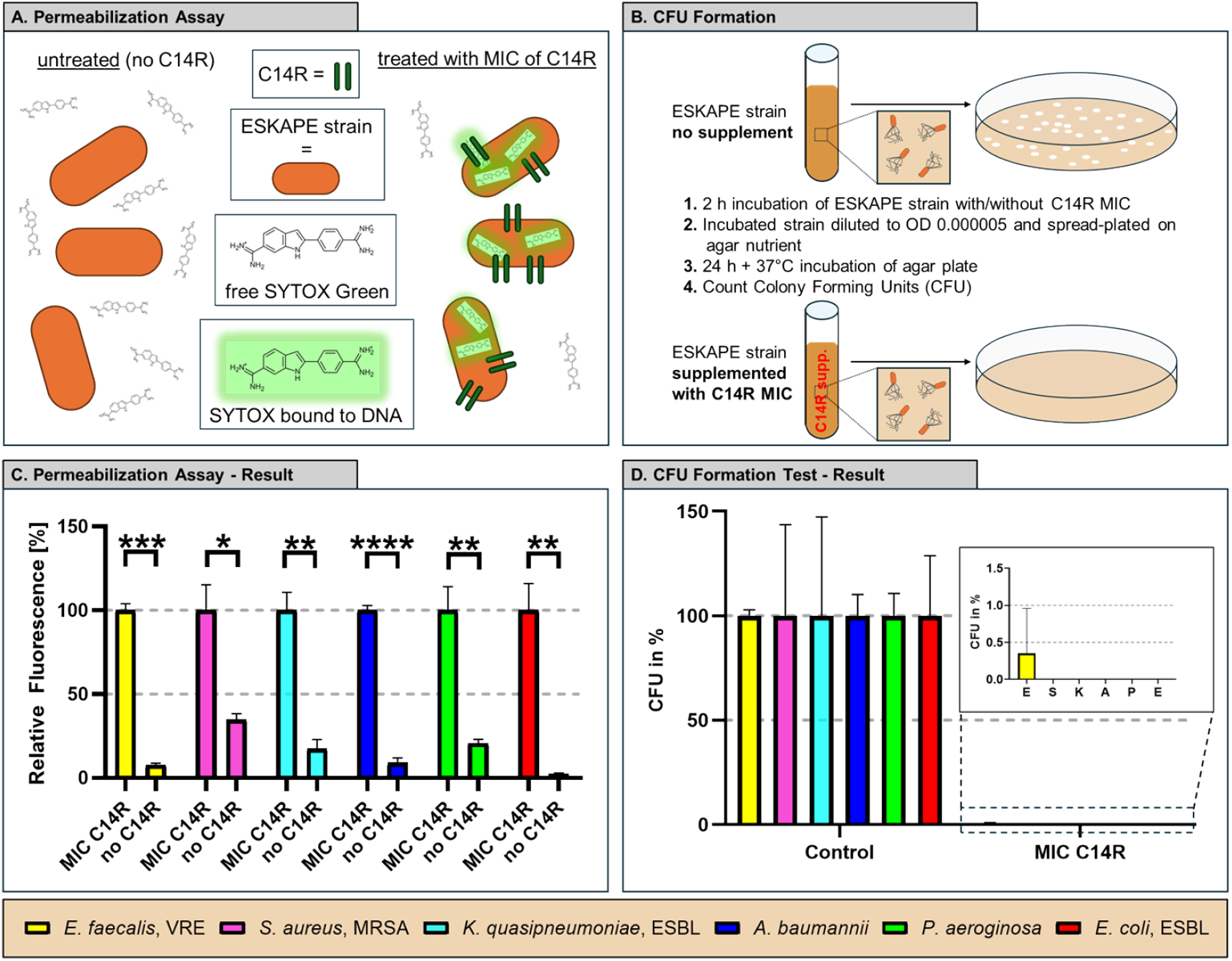
C14R MIC exposure strongly induces membrane permeabilization and reduces bacterial viability (CFU) in all ESKAPE pathogens. (A) Schematic representation of the SYTOX Green membrane permeabilization assay. ESKAPE strains were incubated with C14R, followed by addition of SYTOX Green dye, which selectively enters membrane-compromised cells and fluorescence upon binding to DNA. (B) Colony-forming unit (CFU) assay setup. ESKAPE cultures were exposed for 25 h either to medium alone (control) or C14R peptide at MIC concentrations, followed by plating the incubated cell on agar nutrient plates. After 24 h incubation of the plates, CFUs were counted on control compared to C14R pretreated cells. (C) Membrane permeabilization of ESKAPE pathogens treated with the determined strain-specific C14R MIC. SYTOX Green uptake was measured afterwards, given that increased fluorescence indicates compromised membrane integrity. (D) Results of CFU formation testing. All experiments were done in triplicates. Statistical analysis was performed using the unpaired Student’s t-test. Statistical significance: \**p* < 0.05, \*\**p* < 0.01, \*\*\**p* < 0.001, \*\*\*\**p* < 0.0001. Error bars represent the mean +/-SD.

## Discussion

C14R displayed robust activity against all six ESKAPE pathogens across different tests, confirming a broad antibacterial spectrum. These findings are consistent with earlier C14R studies that classified the peptide as pore-forming and membrane-targeting, with potent activity against *P. aeruginosa* and low cytotoxicity towards lung fibroblasts (24). Beyond antibacterial activity, C14R has been shown to act against fungal pathogens, including *Candida albicans, Candidozyma auris* (formerly *Candida auris)* and *Candida parapsilosis (23)(25)*. Likewise, C14R exhibits antifungal activity against additional clinically relevant species from the WHO fungal priority list, such as *Cryptococcus neoformans, Cryptococcus gattii, Nakaseomyces glabratus, Candida tropicalis, Pichia kudriavseveii* and *Candida dubliniensis*, further underscoring its broad-spectrum antimicrobial potential (28). The cross-kingdom activity of C14R, spanning major ESKAPE bacteria and clinically important fungi like all *Candida* species, suggest application niches where mixed infections are common (e.g. device-associated infections, chronic wounds, blood stream infections). The limited antibiofilm effect on several Gram-negative bacteria seen here argues for combination strategies (with matrix-disrupting enzymes, chelators or antibiotics) and delivery formats (hydrogels) that increase local concentration and residence time. Such formats have been investigated as an antimicrobial drug layer within a two-layer hydrogel, where the upper layer serves as a trapping zone, for example through the incorporation of high affinity-aptamer binding entities against *P. aeruginosa* (29). In this context of rising antimicrobial resistance among high-risk pathogens, the broad-spectrum and pore-forming activity of C14R highlights its promise as a therapeutic candidate to extend current antimicrobial treatment options.

## Material and Methods

### Bacterial strains and cultivation

Liquid cultures of *E. coli, A. baumannii* and *P. aeruginosa* were grown in lysogeny-broth (LB-Miller) overnight at 37 °C with shaking (160 rpm). Other strains were cultured in Todd–Hewitt-broth (Oxoid) supplemented with 0.5% yeast extract (BD, USA). All strains were obtained from Institute of Medical Microbiology and Hygiene, University Clinic of Ulm. (*Enterococcus faecium* (DSM 17050), *Staphylococcus aureus* (ATCC 43300), *Klebsiella quasipneumoniae* (ATCC 700603), *Acinetobacter baumannii* (ATCC 19606), *Pseudomonas aeruginosa* PAO1 and *Escherichia coli* (BSU1286). Cultivation of these strains was carried out as described by Bauer R. et al (30). *K. quasipneumoniae* recently got reclassified from *K. pneumoniae* to *K. quasipneumoniae* by American Type Culture Collection (ATCC).

### Radial Diffusion Assay

Antimicrobial activity was assessed using an overlay radial diffusion assay, as previously described (31). Briefly, target bacteria were inoculated into liquid agarose at a density of 2 × 10^7^ cells per plate. Cultures were inoculated with 1 mL of washed overnight cultures. Wells, put into the solidified plate, were filled with C14R at concentrations ranging from 3.125 to 100 μg/mL. After 3 h incubation at 37 °C, plates were overlaid with tryptic soy agar and incubated overnight at 37 °C in 5% CO_2_. Inhibition zones were then measured.

### Viability/Resazurin Assay

The viability of the ESKAPE pathogens in the presence of C14R (0-200 μg/mL) was assessed according to the CLSI guidelines for antimicrobial susceptibility testing (M100) (27) using Mueller-Hinton-Broth (MHB). Bacterial cultures were adjusted to OD_600_ = 0.0001 and inoculated into 200 μL MHB supplemented with C14R in flat-bottomed 96-well plates, followed by incubation at 37°C for 24 h and 900 rpm. Viability was determined using a modified procedure apart from the CSLI guidelines using a resazurin reduction assay, in which 20 μL of a 0.15 mg/mL resazurin solution was added and incubated for 2 h at 37°C. Conversion of resazurin to resorufin was quantified fluorometrically (λ_Ex_ = 535 nm; λ_Em_ = 595 nm) using a Tecan Infinite F200 microplate reader (Tecan Group Ltd., Männedorf, Switzerland). (32)(33) Minimum inhibitory concentrations (MICs) of C14R were determined via Gompertz fit analysis (data provided in repository). All experiments were done in triplicates.

### Biofilm/Crystal-Violet Assay

Biofilm formation of ESKAPE pathogens was analyzed using the crystal violet assay originally established for bacterial biofilms by George O’Toole (34). To start an initial inoculum of OD_600_ = 0.0001 in Todd-Hewitt-Broth (*E. faecium, S. aureus, K. quasipneumoniae*) and LB medium (*A. baumannii, E. coli, P. aeruginosa*) was prepared (35). Biofilms were exposed to different concentrations of C14R (0-200 μg/mL) diluted in phosphate-buffered saline (PBS) (Life Technologies, Carlsbad, CA, USA). All experiments were done in triplicates.

### Colony Forming Unit (CFU) Assay

Overnight cultures were adjusted to OD_600_ = 0.0001 and incubated for 2 h at 37°C either without peptide (control) or with the strain-specific MIC concentration of C14R (Table 1) in 500 μL total volume. After 2 h incubation 100 μL of the incubated cells were plated on agar, to prevent a bacterial lawn and incubated for 24 h at 37°C. All experiments were done in triplicates.

### SYTOX Green Membrane Permeabilization Assay

Membrane permeabilization was analyzed using the SYTOX Green assay. Overnight cultures were adjusted to OD_600_ = 0.1, centrifuged (4000 x g, 5 min) and resuspended in 10 mM PBS containing 0.5 μM SYTOX Green (Invitrogen). 90 μL of cells in SYTOX Green solution were mixed with PBS only and C14R in PBS at strain-specific MIC. After 10 min incubation, fluorescence (excitation 488nm/emission 530 nm) was measured using a Tecan Infinite M200 plate reader. Longer C14R incubation did not show higher fluorescence values. All measurements were performed in triplicates.

## Funding

This work was supported by the Gesellschaft für Forschungsförderung (GFF) of Lower Austria as part of the project “Aptamers and Odorant Binding Proteins – Innovative Receptors for Electronic Small Ligand Sensing” (FTI22-G-012) and the Förderstelle Wirtschaft, Tourismus und Technologie (WST) (WST-F-5035462/004-2024). This work was also sup-ported by the Austrian Research Promotion Agency (FFG) within the COMET Project “PI-SENS” (project no. 915477) as well as by the Federal Provinces of Lower Austria and Tirol. It was also supported by the Deutsche Forschungsgemeinschaft (DFG, German Research Foundation) project 465229237 and the Anschubfinanzierung A 2025 of Ulm University.

## Data availability statement

Data available at: https://data.mendeley.com/datasets/64vwvnpchv/1

## Notes

### Competing Interest Statement

The authors have declared no competing interest.

